# Genome and community-level interaction insights on wide carbon utilizing and element cycling function of Hydrothermarchaeota from hydrothermal sediment

**DOI:** 10.1101/768564

**Authors:** Zhichao Zhou, Yang Liu, Wei Xu, Jie Pan, Zhu-Hua Luo, Meng Li

## Abstract

Hydrothermal vents release reduced compounds and small organic carbons into surrounding seawaters, providing essential substrates for microbial-derived biosynthesis and bioenergy transformations. Despite the wide distribution of Marine Benthic Group-E archaea (referred to as Hydrothermarchaeota) in hydrothermal environments, little is known on their genome blueprints and ecofunctions. Here, we studied four relatively high-completeness (> 80%) metagenome-assembled genomes (MAGs) from a black smoker chimney and surrounding sulfide sediments in the Mid-Atlantic Ridge of the South Atlantic Ocean (BSmoChi-MAR) as well as publicly available datasets. Comparative genomics suggest that Hydrothermarchaeota members have versatile carbon metabolism, including assimilating proteins, lactate and acetate, degrading aromatics anaerobically, oxidizing C_1_ compounds (CO, formate, and formaldehyde), utilizing methyl-compounds, and incorporating CO_2_ by tetrahydromethanopterin-based Wood–Ljungdahl (WL) pathway and Calvin–Benson–Bassham (CBB) cycle with type III Ribulose-1,5-bisphosphate carboxylase/oxygenase (RubisCO). They could oxidize sulfur, arsenic, and hydrogen, and respire anaerobically via sulfate reduction and denitrification based on genomic evidence. The redundancy of carbon utilizing and element cycling functions, and the interactive processes of syntrophic and sequential utilization of substrates from community-level metabolic prediction, enable wide accessibility of carbon and energy sources to microorganisms. Hydrothermarchaeota members derived important functional components from the community through lateral gene transfer, and became clade-distinctive on genome content, which might serve as a niche-adaptive strategy to metabolize potential heavy metals, C_1_ compounds, and reduced sulfur compounds.

**Importance:** This study provides comprehensive metabolic insights on Hydrothermarchaeota from comparative genomics, evolution and community-level aspects. Hydrothermarchaeota synergistically participates in a wide range of carbon utilizing and element cycling processes with other microbes in the community. We expand the current understanding of community interactions within hydrothermal sediment environments, suggesting that microbial interactions driven by functions are essential to nutrient and element cycling.

## Background

The hydrothermal alterations transfer and deliver reduced sulfur compounds, organic compounds (e.g., C_1_ compounds, petroleum compounds, organic acids, and ammonia) and heavy metals to the surrounding hydrothermal sediments (1-6). Together with deposited sedimentary carbon compounds, these substrates constitute hydrothermal sediments as a distinct ecological niche, compared to deep-ocean cold marine sediments and hydrothermal fluids. In hydrothermal-active Guaymas Basin sediments, microorganisms syntrophically degrade hydrocarbons and lipids, and metabolic linkages among microbial groups were proposed, such as substrate-level interdependency between fermentative members and sulfur- and nitrogen-cycling members (6). However, the diversity and function of hydrothermal environment inhabiting microorganisms, especially archaea, remain elusive and the community-level microbial interactions within these environment settings still lack detailed characterization.

Candidatus Hydrothermarcheota, originally found on continental slope and abyssal sediments and termed Marine Benthic Group E (MBG-E) (7), has recently been proposed as a new archaea phylum (8). A previous study has indicated that Hydrothermarchaeota was an abundant archaea group in the deep-sea hydrothermal environment, such as Juan de Fuca Ridge flank crustal fluids (9). More recently, a study combined the analysis of metagenome-assembled genomes (MAGs) and single-amplified genomes (SAGs) of Hydrothermarchaeota from Juan de Fuca Ridge flank crustal fluids to evaluate the evolutionary placement and functional potential of this new archaea phylum (8), which suggests a potential for carboxydotrophy, sulfate and nitrate reduction of Hydrothermarchaeota (8). Additionally, another metabolic potential analysis based on Hydrothermarchaeota genomes reconstruction from metagenome of Southern Mariana Trough sulfide deposits also suggests their carboxydotrophic and hydrogenotrophic lifestyle (10). However, a relatively small number of available genomes have limited our understanding of the ecological roles and metabolisms of this widely distributed archaeal lineages.

Here, we analyzed metagenomes from sulfur-rich hydrothermal sediments at an active deep-sea (2,770 m depth) hydrothermal vent site (black smoker) in the southern Mid-Atlantic Ridge of the South Atlantic Ocean. (Total S = ∼100-450 mg/g, detailed sample information refers to Supplementary Information). We obtained two metagenomic libraries from the layer (TVG10) and surrounding sediments (TVG13) of an active black smoker chimney in the Mid-Atlantic Ridge (BSmoChi-MAR) of South Atlantic Ocean (38.1 gigabases for TVG10 and 30.3 gigabases for TVG13). *De novo* metagenome assembling and binning resulted in 140 MAGs (> 50% genome completeness) from 24 microbial groups (Table S2), including 5 archaeal MAGs and 135 bacterial MAGs. The metabolic prediction from all resolved MAGs reveals the functional redundancy and syntrophic substrate-utilizing interactions among microorganisms. As implicated from the four relatively high-completeness (> 80%) Hydrothermarchaeota genomes from this study, results of a previous publication (9) and publicly available datasets, we are developing a metabolic scheme of this widely distributed sedimentary archaeal lineage. Evolutionary analysis suggests the important role of lateral gene transfer in the niche-adaptation of Hydrothermarchaeota to surrounding environments. This study provides an advanced insight into the genomics, community-level interactions, and evolution of Hydrothermarchaeota.

## Results and discussion

### Hydrothermarchaeota as a novel archaeal phylum

Reconstructed archaeal MAGs and scaffolds containing phylogenetically informative genes, at least 3 ribosomal proteins (RPs) or 16S rRNA gene fragments, are summarized in Table 1 and Table S3. Both 16S rRNA and RP phylogenies place Hydrothermarchaeota as a distinct lineage parallel to other Euryarchaeotal clades, including Thermococci, Methanomicrobia, and Hadesarchaea (Fig. 1 and Fig. S1). Within this lineage, the 16S rRNA gene sequences show median sequence identities of 80.8-83.9 %, which supports phylum-level diversity (Fig. 1 and Table S4). The phylum designation Hydrothermarchaeota was proposed since all current genomes were obtained from hydrothermal sediments or fluids (7-9). However, 16S rRNA gene sequence data show that Hydrothermarchaeota also occur widely in estuarine and marine sediments, wetland and hot spring sediments (Fig. 1).

**Table 1.** Overview of genomic statistics of archaeal MAGs constructed from this study, the reference, and NCBI-SRA deposits.

| MAGs | Hydrotherm.<br>SZUA-158<br>(TVG10) | Hydrotherm.<br>SZUA-236<br>(TVG13) | Hydrotherm.<br>SZUA-237<br>(TVG13) | Hydrotherm.<br>JdFR-16 | Hydrotherm.<br>JdFR-17 | Hydrotherm.<br>JdFR-18 | Hydrotherm<br>HyVt-292 |
| --- | --- | --- | --- | --- | --- | --- | --- |
| Copies of individual markers |  |  |  |  |  |  |  |
| 0 | 9 | 21 | 49 | 99 | 87 | 2 | 30 |
| 1 | 139 | 128 | 136 | 25 | 59 | 145 | 141 |
| 2 | 1 | 0 | 3 | 25 | 29 | 2 | 17 |
| 3 | 0 | 0 | 0 | 0 | 12 | 0 | 0 |
| 4 | 0 | 0 | 0 | 0 | 1 | 0 | 0 |
| 5+ | 0 | 0 | 0 | 0 | 0 | 0 | 0 |
| Completeness (%) | 92.53 | 84.97 | 65.82 | 31.93 | 53.87 | 98.13 | 80.43 |
| Contamination (%) | 0.93 | 0.00 | 1.76 | 13.55 | 25.15 | 1.87 | 9.06 |
| Strain heterogeneity (%) | 0.00 | 0.00 | 33.33 | 84.00 | 71.83 | 0.00 | 35.29 |
| Bin size (bp) | 1749358 | 1270198 | 1228833 | 1353114 | 2178134 | 2062134 | 1426722 |
| N50 (bp) | 9590 | 6219 | 3825 | 6267 | 7687 | 149032 | 5600 |
| Mean scaffold length (bp) | 7706.42 | 5747.50 | 3510.95 | 5614.58 | 6331.78 | 93733.36 | 4988.54 |
| GC (%) | 38.99 | 47.58 | 45.76 | 49.68 | 49.64 | 39.14 | 40.02 |
| GC standard deviation (%) | 1.91 | 1.99 | 2.89 | 3.92 | 4.50 | 1.54 | 1.98 |
| Coding density (%) | 84.96 | 91.48 | 84.96 | 91.98 | 91.94 | 92.12 | 89.88 |

“Hydrotherm.” stands for Hydrothermarchaeota. HyVt-292 was reconstructed from a metagenome of deep-sea massive sulfide deposits from Southern Mariana Trough (DRR093004), which is hydrothermally inactive. JdFR MAGs were reconstructed from Juan de Fuca Ridge flank subsurface fluids.

**Figure 1.**
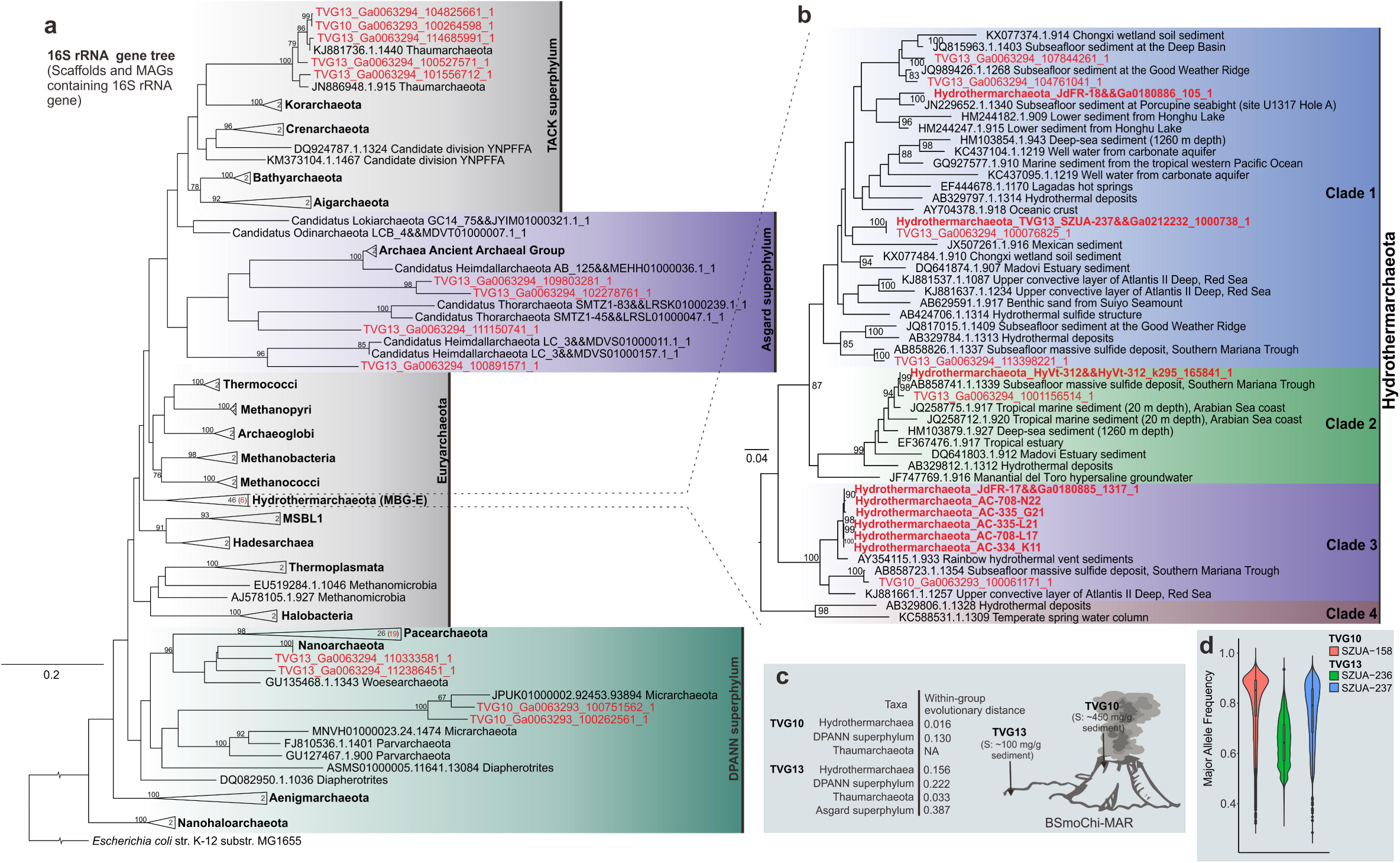
Phylogenetic tree of 16S rRNA gene from TVG metagenomes. **a**, The RAxML maximum likelihood tree was constructed by including metagenome 16S gene sequences (from both binned and unbinned scaffolds). Sequences from this study are highlighted; the highlighted number in bracket stands for sequences from this study. Bootstrap values over 75% were labeled. This tree was rooted by an *Escherichia coli* K12 16S rRNA gene. **b**, Detailed Hydrothermarchaeota 16S rRNA gene tree. Four clades were established based on 90% similarity cutoff 16S rRNA gene representative sequences of Hydrothermarchaeota. Sequences from this study are highlighted. **c**, Schematic figure depicting two sampling locations and with-in group evolutionary distances of 16S rRNA gene sequences from two samples (calculated by Jukes-Cantor model and pairwise comparison based on global alignment). “NA” means only one sequence within a group, not applicable to calculate the with-in group evolutionary distance. **d**, The major allele frequency diagram representing the diverse level of potential Hydrothermarchaeota in corresponding environments where the Hydrothermarchaeota MAGs were found. Higher frequency indicates a majority allele is dominant over the minor ones. Higher mean major allele frequency on the genome level indicates a less diverse Hydrothermarchaeota population in the environment.

### Mixotrophic lifestyle and versatile substrate utilization

We picked four representative Hydrothermarchaeota MAGs of relatively high completeness values (> 80%) from the major three clades for metabolic prediction analysis (Table 1, Fig. 2, Fig. S3 and Tables S5, S6, S7). Similar with the previous study (8), almost all Hydrothermarchaeota MAGs contain the THMPT (tetrahydromethanopterin) based Wood–Ljungdahl pathway (THMPT-WL pathway), and some components of THF (tetrahydrofolate) based Wood–Ljungdahl pathway (THF-WL pathway) (Fig. 2). Since none of them contains the complete genes for THF-WL pathway, it might be that this pathway is not active in Hydrothermarchaeota (Fig. 2). THMPT-WL pathway in Hydrothermarchaeota could function in both directions, either reductively incorporating CO_2_ into acetyl-CoA synthesis or oxidatively converting products from central carbon metabolism (peptide and sugar carbohydrate degradation) into energy producing pathways. If the former direction is active, Hydrothermarchaeota probably lives a mixotrophic lifestyle on using both inorganic and organic carbon sources. Hydrothermarchaeota does not have the methyl coenzyme M reductase (MCR) for methane metabolism, but JdFR-18 could incorporate a variety of methyl-containing compounds into WL pathway, including mono-/di-/trimethylamine and methanol, which is frequently discovered in members of Methanosarcinales, Methanomassiliicoccales, Methanofastidiosa, Bathyarchaeota and Verstraetearchaeota (11). Particularly, JdFR-18 (Clade 1) contains HdrD (3 copies) and GlcD (4 copies, FAD-containing dehydrogenase, similar to D-lactate dehydrogenase) with one pair of them collocated, which is responsible for heterodisulfide reduction linked to lactate utilization. This gene arrangement and function is also present in *Archaeoglobus fulgidis*, Bathyarchaeota, and Verstraetearchaeota (11-13).

**Figure 2.**
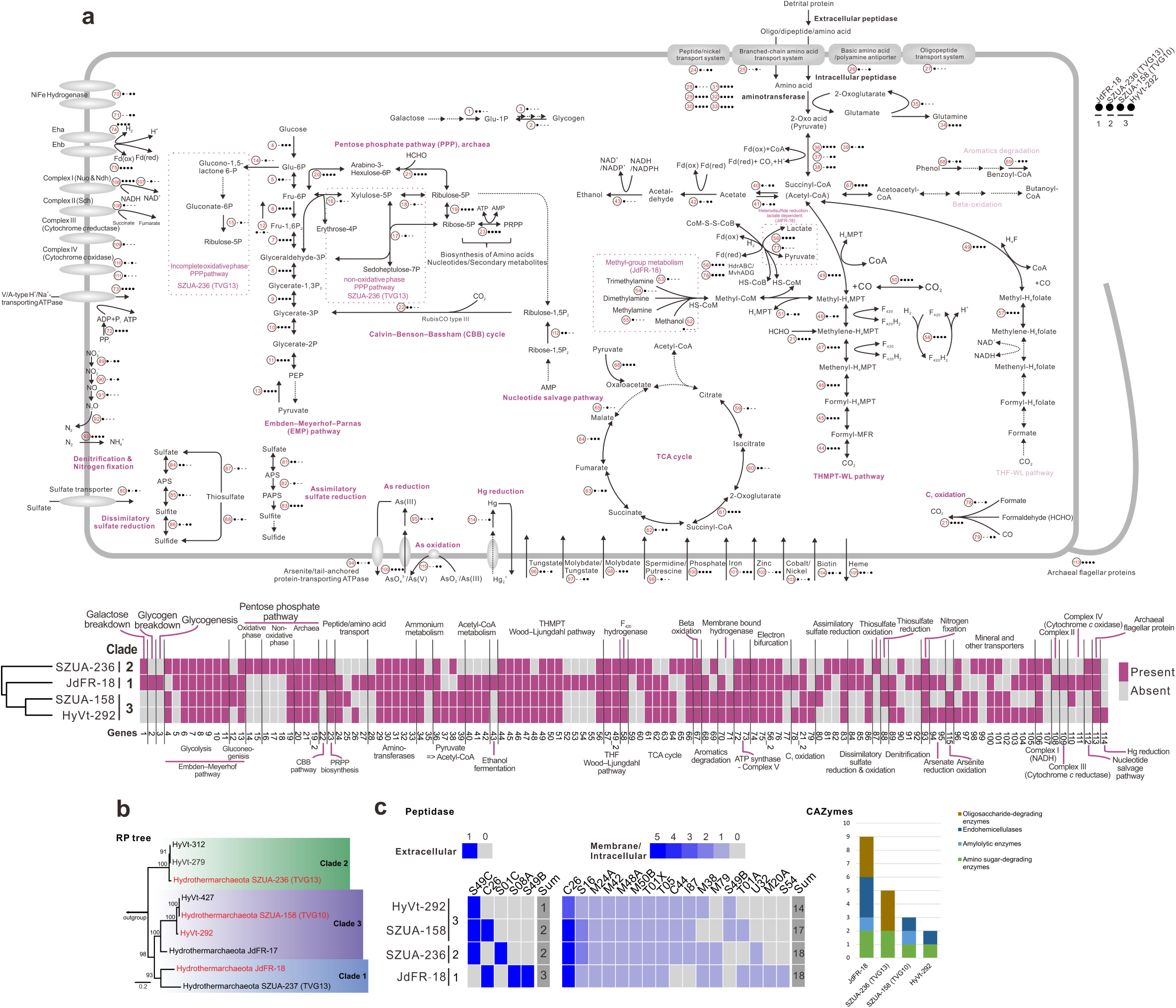
Metabolic pathways of Hydrothermarchaeota. **a**, Schematic figure showing metabolisms of four high completeness Hydrothermarchaeota MAGs. **b**, RP tree showing the phylogeny of all currently available Hydrothermarchaeota MAGs. **c**, Heatmap and barchart depicting the distribution of extracellular/intracellular peptidases and carbohydrate-active enzymes (CAZys) in Hydrothermarchaeota MAGs. The barchart shows the summary information of glycoside hydrolases (GH) family within each functional category.

Hydrothermarchaeota genomes probably encode full TCA cycle, but not beta-oxidation pathway (only acquiring acetyl-CoA C-acetyltransferase coding genes, Fig. 2). Benzoyl-CoA reductase subunits (BcrBC) encoding genes are present in HyVt-292 and JdFR-18, indicating the presence of ATP-dependent benzoyl-CoA degradation. Benzoyl-CoA is the central intermediate in anaerobic degrading pathways of many aromatics, including benzene, phenol, 4-OH-benzoate, cresols, phenylacetate, ethylbenzene and etc. (14). Three out of four Hydrothermarchaeota MAGs contain vanillate/4-hydroxybenzoate decarboxylase subunit C (BsdC) and flavin prenyltransferase (UbiX), further supporting that phenol degradation is possible for them. The existence of both ADP-forming acetyl-CoA synthetase (Acd, EC:6.2.1.13) and acetyl-CoA synthetase (Acs, EC:6.2.1.1) suggests the feasibility of both acetate fermentation and acetogenesis. Further fermentation from acetaldehyde (by Aor) to ethanol (by alcohol dehydrogenase, AdhP/AdhE) has also been discovered in some Hydrothermarchaeota MAGs (Fig. 2).

### C_1_ oxidation and other element cycling capacities

Hydrothermarchaeota could anaerobically oxidize CO for ferredoxin generation, with the existence of carbon monoxide dehydrogenase catalytic subunit (CooS) (Clade 3) (8, 15). NAD^+^-dependent formate dehydrogenase operon of fdsABG exists in both JdFR-18 and HyVt-292, indicating Hydrothermarchaeota could use formate as bioenergy source for electron-transferring phosphorylation (16). The fused 3-hexulose-6-phosphate synthase/6-phospho-3-hexuloisomerase (Hps-Phi) and bifunctional enzyme Fae-Hps are responsible for formaldehyde fixation in ribulose-monophosphate cycle (part of PPP pathway) and generation of methylene-H_4_MPT (THMPT-WL pathway) (EC:4.2.1.147, 4.1.2.43) (3), which are responsible for further biosynthesis of generating ribose and acetyl-CoA, respectively. C_1_ compounds of various redox states are common (CO, formate) or potentially available (formaldehyde) through geochemical reactions in hydrothermal environments (1-4). Additionally, CO could also be generated by some anaerobes (4). Combined with CO_2_ fixation ability as described above (CBB cycle and WL pathway), the mixotrophic lifestyle possibly makes Hydrothermarchaeota as one of the successful archaeal lineages within the global benthic environmental settings (17).

Additionally, the process of sulfide oxidation to sulfate might be possible in Hydrothermarchaeota because the encoded dissimilatory sulfite reductase (DsrAB) could also convey sulfide oxidation (8, 18). The existence of key genes in Sox pathway in SZUA-236 (TVG13) suggests that they could also oxidize thiosulfate for energy yield (Fig. 2). The potential denitrification and sulfate reduction are enable Hydrothermarchaeota to scavenge diverse organic matters by anaerobic respiration. Presumably, Hydrothermarchaeota could couple nitrate reduction with reduced sulfur compound (S^0^, S^2-^ and S_2_ O_3_^2-^) oxidation as the energy generating process (19). As heavy metals are commonly enriched in the hydrothermal environments (20), Hydrothermarchaeota also acquires genomic components for detoxicating As (V) [arsenate reductase (ArsC) and arsenite/tail-anchored protein-transporting ATPase (ArsA)] and Hg (II) [mercuric reductase (MerA)]. Meanwhile, they could also oxidize As (III) [cytomembrane-bound arsenite oxidase subunits (AioB)] (Fig. S2) and presumably couple the reduction of As (V) with the oxidation of reduced sulfur compounds, suggesting that the As cycling could be one of their energy metabolisms.

### Co-existence of nucleotide salvage pathway and CBB cycle

Almost all members from Hydrothermarchaeota encode Embden–Meyerhof–Parnas pathway (EMP pathway) in both glycolysis and gluconeogenesis directions [almost all contain fructose-1,6-bisphosphatase (FBP) and phosphoenolpyruvate synthase (PEP synthase)/pyruvate phosphate dikinase (PPDK)] (Fig. 2). Within glycolysis direction, the conversion of PEP to pyruvate (catalyzed by pyruvate kinase) is lacking in all genomes, however, reverse reactions of PEP synthase/PPDK are reported in some thermophilic archaea, including, *Thermococcus* (Euryarchaeota) and *Thermoproteus* (Crenarchaeota) (21). All Hydrothermarchaeota clades contain archaeal style pentose phosphate pathway (PPP pathway), while, besides that, SZUA-236 (TVG13) contains an incomplete oxidative phase PPP pathway and a non-oxidative phase PPP pathway. The PPP pathway together with phosphoribosyl pyrophosphate (PRPP) synthesis pathway are anabolic for biosynthesis of a variety of amino acids, nucleotides and other secondary metabolites, using substrates from glycolysis (22). They probably fix CO_2_ by type III-RubisCO in the Calvin–Benson–Bassham (CBB) cycle based on genomic prediction (Fig. 2). The lacking of phosphoribulokinase (Prk) of CBB cycle is frequently seen in archaeal genomes (23); meanwhile, some reports based on metabolic experiments indicate the presence of autotrophic activity of crenarchaeotal CBB cycles despite lacking Prk, suggesting potential existence of its function in Hydrothermarchaeota (24). The existence of nucleotide salvage pathway and CBB cycle suggests that Hydrothermarchaeota could recover the RNA/DNA degradation products (adenosine monophosphate, AMP) into glycolysis or cycle them back into PPP pathway for biosynthesis (Fig. 2) (25). Beside the RNA/DNA degradation, AMP could also be originated from activities of i) AMP-forming adenylylsulfate reductase during sulfate reduction (Clade 1, 2), ii) PRPP synthesis process (all clades), iii) ADP-dependent (AMP-forming) phosphofructokinase/glucokinase (Clade 3) during glycolysis (26-28). The genomic components of type III-RubisCO and nucleotide salvage function are currently found in other euryarchaeotal groups, including, Archaeoglobi, Halobacteria, Thermococci, Hadesarchaea, and euryarchaeotal methanogens (1). Unconventional participation of type III-RubisCO in nucleotide salvage function suggests the primary function of type III-RubisCO in early ages of archaea evolution (1, 25).

### Limited carbohydrate assimilation while being protein/peptide degraders

No potential sugar and carbohydrate transporters have been discovered among the major three clades of Hydrothermarchaeota (Fig. 2) and they encode limited functions of carbohydrate assimilation and transformation, only including galactose degradation and glycogen conversion. The annotation of CAZys also suggests they have limited capacity in carbohydrate utilization, among which serine-type endopeptidase S08A is the dominant extracellular peptidase from both metagenome and metatranscriptome of MG-I, -II and -III archaea of deep-sea hydrothermal plume (29); S49C is the archaeal signal peptide peptidase for destructing cleaved signal peptides; C26 is the gamma-glutamyl hydrolase and probably acquires the glutamine amidotransferase activity (Tables S5, S6, S7). Meanwhile, genomic predictions indicate that Hydrothermarchaeota (from Clade 1 and 3) have various peptide/amino acid transporters, and all the major three clades acquire six groups of aminotransferases for transferring amino residues (Tables S5, S6, S7), and pyruvate ferredoxin oxidoreductase (Por), indolepyruvate ferredoxin oxidoreductase (Ior), 2-oxoglutarate/2-oxoacid ferredoxin oxidoreductase (Kor) and pyruvate dehydrogenase, dihydrolipoamide dehydrogenase for assimilating 2-oxo acids (pyruvate) to succinyl-CoA (acetyl-CoA) and replenishing the energy pool of reducing equivalents. The existence of encoded proteins of various peptide/amino acid transporters and endopeptidases/aminotransferases suggests that Hydrothermarchaeota use detrital peptides/proteins as one of the main carbon and energy sources.

### Function redundancy and community level interactions

We have analyzed the metabolic capacities of all reconstructed MAGs (Fig. 3 and Tables S2, S8) to investigate microbial community interactions. Acidobacteria, Bacteroidetes, and Gemmatimonadetes acquire the most abundant genes encoding for extracellular peptidases; they are presumably the major players in utilizing detrital proteins from marine sediments. Other microbial groups could form syntrophic interactions with them for assimilating extracellular peptides/proteins using the extracellular peptidases secreted by them. Furthermore, Ignavibacteriae, Planctomycetes, and Spirochaetes acquire the most abundant genes encoding for glycoside hydrolases, suggesting that they are the major players in carbohydrate/sugar utilization. Besides, a variety of microbial groups are predicted to acquire degrading/utilizing ability on methane, fatty acids, aromatics, methanol, and mono-/di-/trimethylamine. The fermentation products probably include acetate, hydrogen, lactate, and ethanol. The electron pools generated through the fermentation steps are delivered to terminal electron acceptors or CO_2_ for either respiration or C fixation. Moreover, the fermentation products from the first fermentation process could also be re-utilized by the community members as energy and carbon sources. The interaction among microorganisms of syntrophic and sequential (step by step) utilization of substrates enables the community to gain more energy from a wide range of substrates.

**Figure 3.**
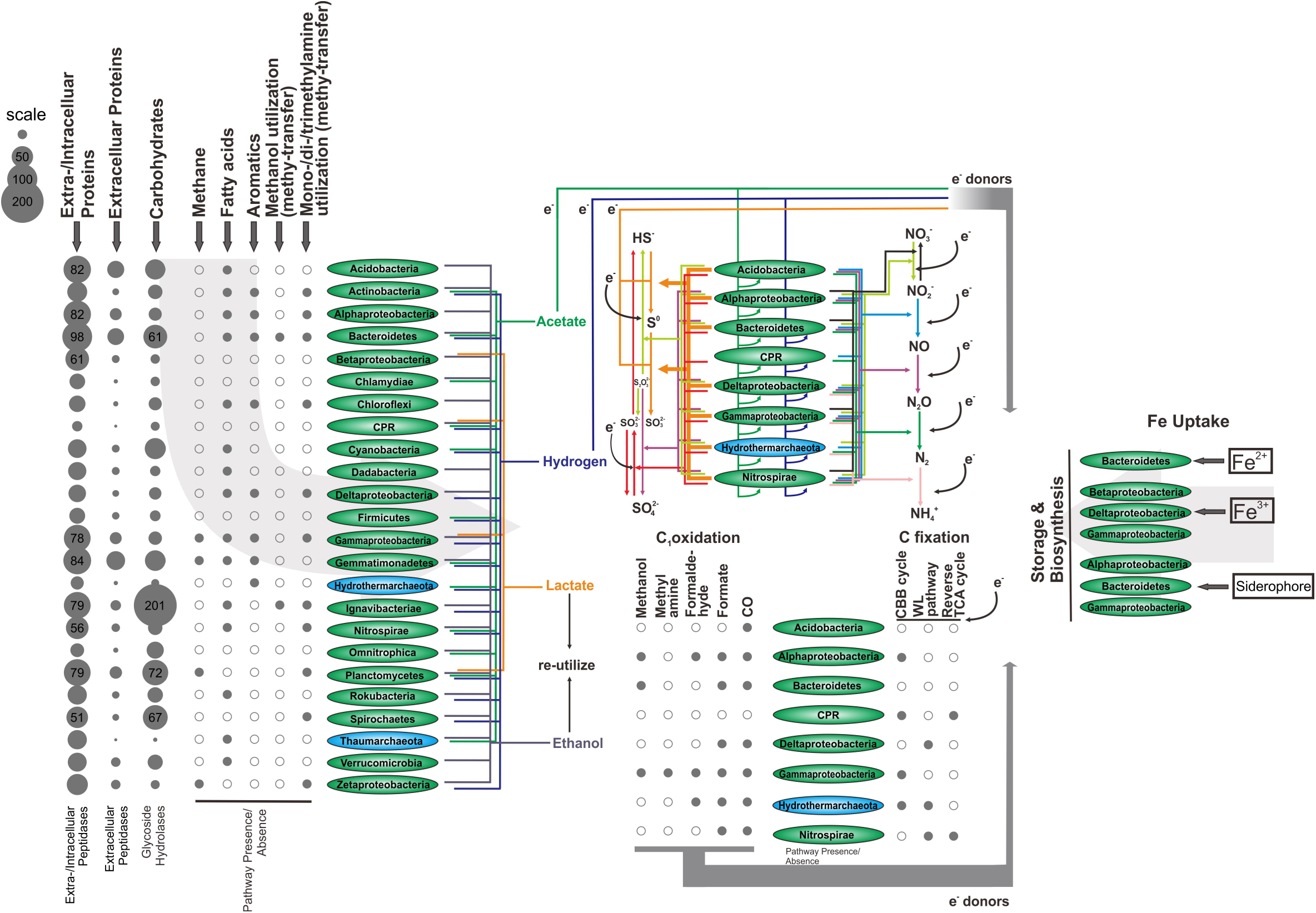
Metabolic prediction figure for the community of MAGs from TVG metagenomes. The peptidases and carbohydrate degrading enzymes were calculated by counting the MAG completeness and taking average values of all MAGs within individual microbial groups (0 digits after the decimal point). The presence of specific pathway/function within each microbial group was assigned when this pathway/function was present in any MAGs within this microbial group. The Fe uptake metabolism was predicted by the corresponding database. For the metabolic prediction of N, S cycling, C_1_ oxidation, and C fixation, only the major microbial groups (with at least one MAG from this group acquiring genome coverage > 15×) are represented.

The major eight microbial groups (with at least one MAG from this group acquiring genome coverage > 15×, including Acidobacteria, Alphaproteobacteria, Bacteroidetes, Candidate Phyla Radiation, Deltaproteobacteria, Gammaproteobacteria, Hydrothermarchaeota, Nitrospirae) are predicted to acquire multiple functions on sulfur cycling, including sulfide oxidation, sulfur oxidation, thiosulfate oxidation, and sulfate reduction, thiosulfate disproportionation. The oxidized sulfur compounds (SO_4_^2-^, SO_3_^2-^ and S_2_O_3_^2-^), as well as nitrate/nitrite and molecular oxygen [except for Candidate Phyla Radiation (CPR) and Hydrothermarchaeota], could serve as the terminal electron acceptors for respiration on organic or inorganic energy sources (Table S8). This suggests that microorganisms in the chimney layers and surrounding sediments of BSmoChi-MAR acquire various strategies adapting to microaerobic to anoxic environment settings. Besides of Hydrothermarchaeota, several other microbial groups are also predicted to oxidize multiple C_1_ compounds and acquire C fixation capacity in their genome contents; in addition, some of these microbial groups are probably capable for carbohydrate and peptide/protein degradation and acquire sulfur cycling and denitrification abilities, such as Alpha-/Delta-/Gammaproteobacteria and Nitrospirae (Fig. 3 and Table S8). It is suggested that the redundancy of carbon utilizing and element cycling functions of microorganisms and the interactive processes of syntrophic and sequential (step by step) utilization of substrates among microorganisms enable a wide range of substrates and energy sources to be accessible to the community.

### Comparative genomics

We chose representative genomes from euryarchaeotal groups and Hydrothermarchaeota to compare the metabolic capacities among them. Peptide degradation capacities are shared among most Hydrothermarchaeota and euryarchaeotal groups, while, the other carbohydrate degrading/utilizing capacities on starch/glycogen, aromatics, fatty acids, methanol, and mono-/di-/trimethylamines are patchily distributed (Fig. 4, Figs. S3, S4 and Table S9). Hydrothermarchaeota acquires considerably complete functions in the cycling of N and S and could oxidize three important C_1_ compounds, comparing to the euryarchaeotal groups. When it comes to within phylum level, we found the distinction of metabolic traits among three major clades within Hydrothermarchaeota (Fig. 4). Clade 1 MAG (from subsurface fluids from SubFlu-JdFR) specifically acquires utilizing ability on methanol, methanethiol, and mono-/di-/trimethylamines, which is not shared with other clades, probably indicating potential supply of methyl-compounds in the surrounding environments. Clade 3 MAGs (SZUA-158 from TVG10, the chimney layer sample, of BSmoChi-MAR and HyVt-292 from sulfide deposits of Southern Mariana Trough) acquire sulfide oxidation ability (DsrAB) as the major sulfur cycling function, probably being attributed to the high supply of sulfide in the surrounding environments. Presumably, Clade 3 could also depend more on sulfide oxidation for energy and acquire less sugar carbohydrate degrading enzymes (Fig. 2). It provides a clue that each clade could acquire metabolic traits related to niche adaptation.

**Figure 4.**
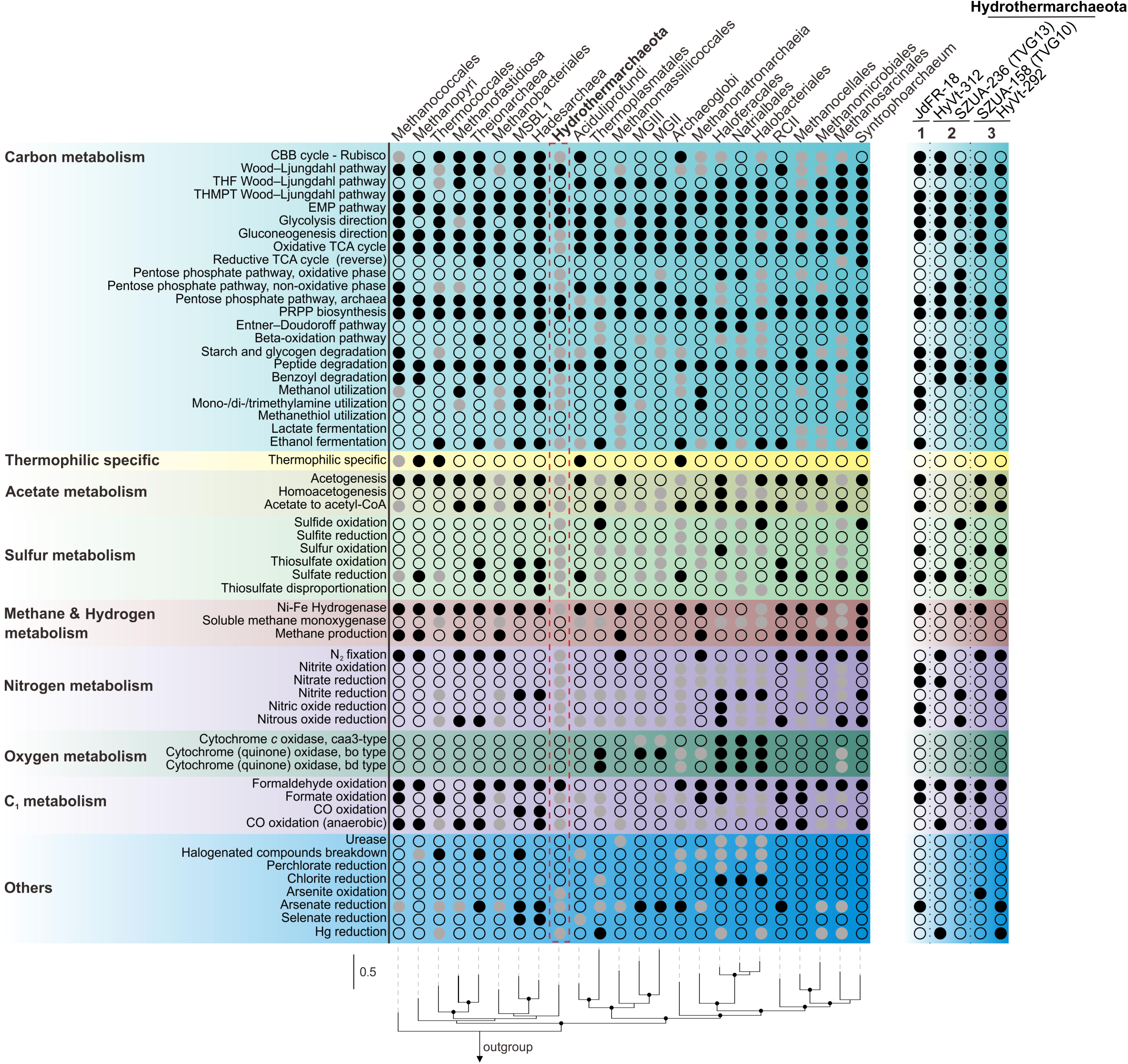
Metabolic capacity comparison between Hydrothermarchaeota and Euryarchaeota. Metabolic marker genes from Pfam, TIGRfam and KEGG databases are used to search for up to five genomes within one archaeal group. Solid black dots, solid grey dots, and blank dots stand for all present, partially present, and no present in all genomes. If one marker gene appears, it is assumed that the corresponding metabolic function exists. Due to the limited genomes and low genome completeness, for Hadesarchaea, Hydrothermarchaeota, Theionarchaea, Syntrophoarchaeum, and MSBL-1, if genes appear in one genome, solid black dots are used. Detailed summary information refers to Supplementary Information. The metabolic capacity of five Hydrothermarchaeota MAGs was also depicted. The RP phylogenetic tree was constructed by picking one random genome from each group and bootstrap values over 75% were depicted as black dots on the node.

### Clade-distinctive lateral gene transfers

We mapped the minimum parsimony-based prediction of gene gain and loss events of inferred gene ortholog groups (OGs) to the RP-based phylogenomic tree (Fig. 5 and Fig. S5). The phylogenomically close-related seven euryarchaeotal classes or orders were included, acquiring both methanogens and non-methanogens. The key gene gain events at node 28, 26 and 24 (occupying 24.9%, 6.7% and 29.1% of the ancestral genomes) indicate that the important traits of extant Hydrothermarchaeota are derived from lateral gene transfers (LGTs); they include the C_1_ oxidation on formaldehyde and CO, the key component of Wood–Ljungdahl pathway for CO_2_ fixation and acetyl-CoA synthesis, nitrogen and sulfur cycling, aromatic degradation and Hg and As reduction. The gene gain events at node 27 (occupying 27.9% of the genome) probably provide JdFR-18 abilities on utilizing tri-/mi-/monomethylamines, acetogenesis and other functions on nitrogen and sulfur cycling (Table S10). The lost OGs at these nodes mainly acquire functions related to amino acid transport and metabolism, energy production and conversion, and transcription and translation related metabolisms (Table S10 and Fig S6). As we have indicated above, it might be the adaptive strategy of Hydrothermarchaeota to derive functional components from the lateral gene interactions among community members in hydrothermal environments, which are characterized with plenty of heavy metals, C_1_ compounds, and reduced sulfur compounds (1-6, 19, 20, 30). The distinctive metabolism of each clade (Figs 4, 5 and Tables S11, S12) on C, H, N, and S could also be due to LGT events in the adapting process within corresponding eco-niches. The OG component pattern tells that although Hydrothermarchaeota could not produce methane, they are close to the anaerobic wastewater treatment inhabiting methanogenic phylum Methanofastidiosa (Fig. S7), which is also consistent to the RP-based phylogenomic tree. Nevertheless, the derived functions through LGT probably make Hydrothermarchaeota distinctive and become the hydrothermal environment-adaptive archaeal lineage.

**Figure 5.**
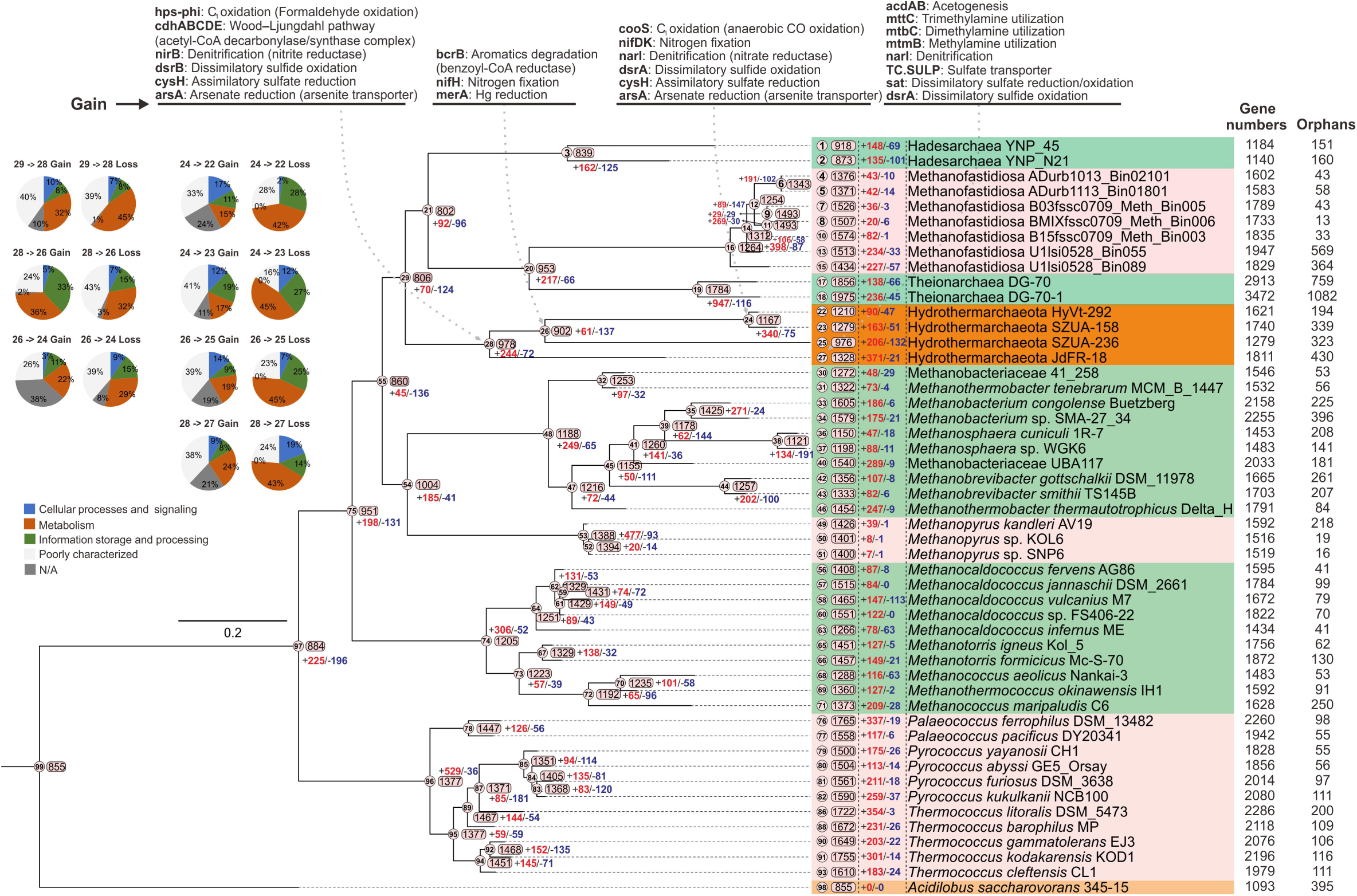
The estimation of ortholog group (OG) turnover events for Hydrothermarchaeota and related euryarchaeotal orders and classes. The OG numbers and inferred OG gain and loss numbers were labeled accordingly on the tree nodes and tips. The COG category information of the gained or lost OGs for Hydrothermarchaeota clade was parsed and depicted. The important genes that were involved with the OG gain events for Hydrothermarchaeota clade were also labeled to the corresponding nodes.

## Conclusions

Hydrothermarchaeota is the widespread archaeal lineage among the archaeal members in the hydrothermal sediment environments. Within the microbial community, Hydrothermarchaeota synergistically participates in a wide range of carbon utilizing and element cycling processes with other microbes. This study suggests that microbial interactions are essential to nutrient and element cycling, and extends the current understanding of community interactions within hydrothermal sediment environments (6, 31). Our findings call for further genomic studies of Hydrothermarchaeota from other environments, including estuarine, wetland and spring sediments, and genomic, transcriptomic and enzymatic studies by cultivation-based experiments to study their metabolic capacities and activities.

## Materials and Methods

### Sample information and metagenome sequencing

Marine hydrothermal sediment samples were retrieved from an active deep-sea hydrothermal vent site (black smoker) of 2,770 m depth in the Mid-Atlantic Ridge of South Atlantic Ocean, during the cruise of DY125-26 by R/V Dayang Yihao (Ocean No. 1) at August 2012 (32). TVG10 was sampled from the layer from a black smoker chimney, and TVG13 was a sulfide sediment sample collected near the black smoker chimney. Samples were stored in −80°C for subsequent metagenome sequencing, and physicochemical characterizations were conducted soon after collection, which included total C, H, S, C/N ratio and pH (32) (Experimental details and results refer to the previous work).

### Metagenome processing and genome-resolved binning

In order to get high quality archaeal metagenome-assembled genomes (MAGs), a custom processing method with two rounds of assembling and binning was adopted. The metagenomes were sequenced by Illumina HiSeq 2000 platform, two separated libraries for each sample were obtained and combined into one in the downstream analysis. Raw reads were firstly dereplicated and processed by Sickle (https://github.com/najoshi/sickle) for trimming reads of low quality with default settings. Clean reads for each sample were subjected to *de novo* metagenome assembly by IDBA-UD v1.1.1 with ‘--mink 52 --maxk 92 --step 8’ settings (33). The initially resulted assemblies were deposited to DOE-JGI IMG (The Integrated Microbial Genomes system of US Department of Energy-Joint Genome Institute) database and annotated by the DOE-JGI Microbial Genome Annotation Pipeline (MGAP v.4) (34).

Assemblies were subjected to a MetaBAT v0.32.4 based binning with 12 combinations of parameters (35), subsequently, Das-Tool v1.0 was applied to screen MetaBAT bins, resulting with high quality and completeness bins (36). CheckM v1.0.7 was used to assess the bin quality and phylogeny (37). All above resulted archaeal MAGs were combined with i) all available archaeal genomes from GenBank database (Aug 2, 2017 updated), ii) archaeal clones, fosmids and cosmids sequences from NCBI Nucleotide database (Aug 2, 2017 updated), iii) initial assembled scaffolds with one or more ORFs annotated as archaeal origin by IMG database (Only assemblies obtained in this study), as the reference for reads mapping. BBmap was used to get potential archaeal reads from raw reads with ‘vslow minid = 0.6’ option (38). The second round of assembling by archaeal reads was the same as the above method, and potential ‘archaea related scaffolds’ were also subjected to DOE-JGI IMG database to get annotated as described above. The same ‘MetaBAT+Das-Tool’ method was used to get the second round of MAGs, and only archaeal MAGs with high quality were used for downstream analysis. Outlier scaffolds with abnormal coverage, tetranucleotide signals and GC pattern within potential high contamination MAGs (checked by CheckM) and erroneous SSU sequences within MAGs were screened out and decontaminated by RefineM v0.0.20 with the default settings (39). Average genome coverages were calculated by remapping raw reads to MAGs using Bowtie2 v2.2.8 (40). The bacterial MAGs were obtained using the similar binning and decontamination processes, but with only one-round binning. Further refinement was also conducted by manual inspection based on VizBin for selective MAGs (41).

SRA information was obtained by searching string “(((hydrothermal) AND metagenomic[Source]) AND WGS[Strategy])) NOT 16S[Title] NOT 454 GS[Text Word] AND (metagenome[Organism] OR hydrothermal vent metagenome[Organism] OR marine sediment metagenome[Organism] OR marine metagenome[Organism] OR subsurface metagenome[Organism])” (Dec 26, 2017 updated) for hydrothermal vent sediment studies and “(((spring sediment) AND metagenomic [Source]) AND WGS[Strategy])) NOT 16S[Title] NOT 454 GS[Text Word]” (Jan 24, 2018 updated) for freshwater spring sediment studies deposited in NCBI-SRA. Searching results were manually inspected to confirm (Table S1). The linked DOE-JGI IMG deposits to these SRA deposits were found and the assemblies are used. MAGs originated from hydrothermal vent sediments (21 studies) and freshwater spring sediments (22 studies) were reconstructed from these public NCBI-SRA deposits and the linked DOE-JGI IMG deposits (Only one study has the IMG record but no SRA record. This study was also manually inspected to meet the searching criterion). SRA runs within one ‘Experiment’ and studies for one ‘Biosample’ are subjected to integrated assembling. Assembling was conducted by MEGAHIT v1.1.2 (42) with kmer iterations of k35-k75, k45-k95, k65-k145, k145k-k295 for 85bp, 100bp, 150bp, and 300bp reads and the kmer step of 10; the pre-processing was the same as described above. Studies which have DOE-JGI IMG deposits were simply used with their assembled metagenomes and QC-passed reads. The downstream binning methods were the same as described above but with one-round binning. Further refinement was also conducted by manual inspection based on VizBin for selective MAGs (41).

### Archaeal MAGs annotation

KO annotation was made by GhostKOALA v2.0, KAAS v2.1 and eggNOG-mapper v4.5.1 (Use its first KO hit and COG hit, COG were translated to KO by ‘ko2cog.xl’ provided by KEGG database) (43-45). Annotation by NCBI nr database (Mar 6, 2017 updated) was done by extracting the first meaningful hit (meaningful information rather than ‘hypothetical proteins’). Peptidases were called by MEROPS (Use its ‘pepunit’ database for less false positive hits) via DIAMOND BLASTP v0.9.10.111 with ‘-k 1 -e 1e-10 --subject-cover 80 --id 50’ settings (46, 47). Carbohydrate-active enzyme (CAZy) annotation was carried out by dbCAN (version 20170913) and interpreted by CAZy database (self-parsed online information) (48, 49). InterProScan 5.26-65.0 (client version) was applied to classify protein functions with annotations including, CDD, PfamA, SMART, TIGRFAM, Phobius, and SuperFamily(50). Phobius, RED-SIGNAL and PSORTb v3.0.2 (Archaea) were applied to predict the location of peptidases, as ‘Membrane/Intracellular’ or ‘Extracellular’ (Only congruent results of ‘Extracellular’ location in all 3 methods were adopted, while, others with incongruent results were assigned as ‘Membrane/Intracellular’) (51-53).

### Major allele frequency analysis

The anvi’o v4.0 was used to identify and profile single-nucleotide variants (SNVs) of Hydrothermarchaeota MAGs based on mapping the reads from corresponding metagenomes. The characterizing strategies for identifying SNVs are operated according to the instruction (http://merenlab.org/2015/07/20/analyzing-variability/). The major allele frequency value was the percentage of metagenomic reads mapping to a certain site with the majority SNV.

### Comparative genomic analysis

The Markov Cluster (MCL) Algorithm implemented in anvi’o v4.0 was applied for protein clustering (54). The eggNOG-mapper v4.5.1 was used to annotate MAGs with default settings (44, 45). COG functional categories and orthologous groups parsed from eggNOG mapping results were used to reconstruct the inner tree. The existence of specific functions or pathways was assigned according to the existence of marker genes (Use the annotation results from the above section). Average nucleotide identity (ANI) values among Hydrothermarchaeota MAGs were calculated by OrthoANI with default settings (55).

### Phylogenetic reconstruction

Searching for sequences in SILVA SSU128 for long Marine Benthic Group E (Hydrothermarchaeota) sequences with good quality (pintail quality > 75%, sequence length > 1000 nt and sequence quality > 75%) resulted in 549 sequences (assigned as Hydrothermarchaeota backbone tree, “HydroBTree”) (56). The obtained alignment was subjected to clustering by mothur (57). 36 OTU representative sequences at 90% similarity cutoff were obtained. Representative sequences in SSURef_NR99_128_SILVA database and archaeal 16S rRNA gene sequences retrieved from metagenomic scaffolds (curated by IMG database) and MAGs were combined (only sequence length > 300bp being considered), and subsequently, subjected to aligning by SINA v1.2.11 (58). The updated 16S rRNA genes from Pacearchaeota and Asgard superphylum genomes (deposited in NCBI Genome database) were also included in the tree construction. The SINA alignment with *Escherichia coli* K12 as the outgroup was filtered by both ssuref:archaea (LTPs128_SSU) and 50% consensus filters, and subsequently used for tree construction by RAxML-HPC v.8 on XSEDE implemented in CIPRES, with settings as GTRCAT and 1000 bootstrap iterations (59, 60).

The 16S rRNA gene sequences (> 300bp) which were BLASTed out from the Hydrothermarchaeota MAGs constructed from NCBI SRAs, the previous publication and this study were aligned by SINA v1.2.11 (58) and inserted into the “HydroBBTree” by “ARB_PARSIMONY quick-add species” method in ARB (61) (Some MAGs have no 16S rRNA gene sequences, which is normal, due to the low MAG completeness). The topology of this 16S rRNA gene tree remains unchanged compared to that of “HydroBBTree” and the division of clades also remains unchanged.

The masked alignment of 12 ribosomal proteins (processed by CheckM, including, L2, L3, L4, L5, L14, L16, L18, L22 and S3, S8, S17, S19 ribosomal proteins) were concatenated and then subjected to the tree model selection by ProtTest 3 (37, 62). Representative archaeal genomes and reported Hydrothermarchaeota MAGs were included in the tree together with MAGs and scaffolds from this study (63). A pre-selection was imposed on the concatenated alignment to filter those sequences with less than 3 ribosomal proteins and less than 25% alignment columns; columns with more than 50% gaps were trimmed. The RAxML-HPC v.8 on XSEDE implemented in CIPRES was applied to make the phylogenetic tree with the best model as PROTGAMMAILG and 1000 bootstrap iterations (59, 60). *Escherichia coli* K12 genome was adopted as the outgroup (64).

### Evolutionary analysis

The genomes from phylogenetically close-related archaeal orders/classes were acquired from NCBI Genome database, including methanogenic Methanobacteriales, Methanococcales, Methanofastidiosa and Methanopyri, and non-methanogenic Theionarchaea, Hadesarchaea, and Thermococcales. One Crenarchaeota (*Acidilobus saccharovorans* str. 345-15) genome was used as the outgroup. The genome picking criterion is that they are over 80% completeness and less than 10% genome contamination, with only exceptions of two Theionarchaea genomes (the only two available genomes within the class) and one Hadesarchaea genome (77.6% completeness; one out of two genomes within the class); and genomes are from different families or genera if possible. The phylogenomic tree of acquired 50 genomes were constructed with the concatenated masked alignment of 12 ribosomal proteins by the same method as described above, but using IQ-TREE v1.6.3 (with better performance) (65) with the settings as “-m MFP -mset LG,WAG -mrate E,I,G,I+G -mfreq FU -bb 1000”.

The ortholog groups (OGs) of protein-encoding genes shared by 50 genomes were parsed out by OrthoFinder v2.2.6 (66) with orphan genes (only existing in one genome) not included in OGs. The BadiRate was used to estimate OG turnover rate using the BDI-FR-CSP model (Turnover rates-Branch model-Estimating procedure, stringent on estimating turnover rates) (67) with the above phylogenomic tree as the input tree file. The output gene turnover results were parsed to OG turnover results by a custom Perl. The OGs were annotated by eggNOG-mapper v4.5.1; each was assigned with the majority annotation result. The key OG turnover events on Hydrothermarchaeota nodes were parsed; the related genes with function and pathway annotations were summarized.

### Metabolic capacity prediction and comparison

Genomes of Euryarchaeota and Hydrothermarchaeota were acquired from NCBI Genome database, and every five representative genomes (picked from different families if possible) from each archaeal group were used (9). Only genomes with completeness over 70% were used. If one archaeal group has limited available genomes (less than five), all the genomes were used, regardless of the completeness. Metabolic marker genes were retrieved from a custom metabolic gene database and metabolic pathways annotated in KEGG database (68, 69). The Pfam, TIGRfam and custom metabolic gene database were used to scan against genomes with suggested cutoff settings; GhostKOALA v2.0, KAAS v2.1, and eggNOG-mapper v4.5.1 were applied to assign KOs to genomes based on default settings (43-45). For each metabolic marker gene, we label their presence/absence in archaeal groups as solid black dots (present in all), solid grey dots (present in some) and blank dots (present in none). For each metabolic function, if one marker gene appears, we assign the presence of this function. Due to the limited genomes and low genome completeness (less than 70%) for Hadesarchaea, Theionarchaea, Syntrophoarchaeum, and MSBL-1, if one metabolic marker gene/metabolic function appears in any genomes within an individual archaeal group, a solid black dot is used.

For the community metabolic analysis on the MAGs from both the metagenomes, the similar metabolic capacity prediction method was used as described above. The peptidases and carbohydrate-active enzymes were calculated by counting the MAG completeness and taking average values of all MAGs within individual microbial groups (0 digits after the decimal point). The presence of specific pathway/function within each microbial group was assigned when this pathway/function was present in any MAGs within this microbial group. The Fe uptake metabolism was predicted by the corresponding database (70), using DIAMOND BLASTP v0.9.10.111 with the settings of ‘-e 1e-20 --query-cover 80 --id 65’ (47).

## Acknowledgments

We thank the cruise of DY125-26 and all crew members who have participated in sampling deep ocean sediments.

## Availability of data

Initial assemblies: IMG: 3300003886 for TVG10 and IMG: 3300003885 for TVG13. Second round assemblies: IMG: 3300020233 for TVG10 and IMG: 3300020236 for TVG13. The MAGs that were resolved from this study are deposited under NCBI BioProject PRJNA385762 and PRJNA480137. The detailed genomic parameters of these assembled genomes are summarized in Supplementary Information.

## Author’s contributions

Z.Z. and M. L. conceived and designed this study. Z.-H. L. and W.X. contributed to sample collection and physicochemical parameter measurement. Z.Z. processed the sequencing data, reconstructed metagenome-resolved genomes and performed the downstream bioinformatic analyses. Y.L., P.J., Z.Z. and M. L. contributed to the bioinformatic infrastructure construction. All the authors were involved in the manuscript writing and approved the final edition of the manuscript.

## Ethics approval and consent to participate

Not applicable.

## Consent for publication

Not applicable.

