## Supplementary material for "Genome and community-level interaction insights on wide carbon utilizing and element cycling function of Hydrothermarchaeota from hydrothermal sediment": Supplementary_Information.docx

**The physicochemical parameters of hydrothermal sediment samples**

| **Sample** | **Depth (m)** | **pH** | **Total C (mg/g)** | **Total N (mg/g)** | **Total H (mg/g)** | **Total S (mg/g)** | **C/N ratio** |
| --- | --- | --- | --- | --- | --- | --- | --- |
| TVG10N | 2770 | 7.09±0.11 | 16.81±0.86 | 2.51±0.61 | 11.03±1.28 | 418.81±10.09 | 7.79±0.76 |
| TVG10W | 2770 | 7.17±0.21 | 15.47±1.36 | 2.45±0.04 | 10.99±0.71 | 480.73±11.33 | 6.32±0.65 |
| TVG13 | 2730 | 7.30±0.27 | 19.16±0.91 | 2.85±0.15 | 10.95±0.41 | 104.72±14.303 | 6.73±0.13 |

The pH was determined in 1:1 sample/water slurries with an acidometer. The other physicochemical characters were analyzed using a Vario EL III Elemental analyzer (Elementar, Germany). The parameter table and analyzing methods are originated from our previous publication^1^. Sample TVG10N and TVG10W were combined to one, as TVG10; Sample TVG10 and TVG13 were used to isolate metagenomic DNA.

### Archaeal genome reconstruction and phylogeny

According to MAGs and informative scaffolds from both RP and 16S rRNA gene trees, TVG10 acquires more Pacearchaeota and DPANN superphylum communities, while TVG13 acquires more Thaumarchaeota and Asgard superphylum communities, respectively (Figure 1 and Supplementary Figure S1). TVG10 acquires low diversity Hydrothermarchaeota MAGs and scaffolds mainly clustered in Clade 3, while, TVG13 acquires high diversity Hydrothermarchaeota MAGs and scaffolds mainly clustered in Clade 1 and 2 (Figure 1b, 1c). There are minor distributions of Asgard and DPANN superphylum in this regime, albeit, with too low genome coverage for reconstructing MAGs (Figure 1). Comparing to TVG10, TVG13 acquires more diverse Hydrothermarchaeota and DPANN superphylum communities as evident from the analysis on within-group 16S rRNA gene evolutionary distance (Figure 1c). The higher mean major allele frequency of SZUA-158 from TVG10 on the genome level also indicates a less diverse Hydrothermarchaeota population in this environment (Figure 1d). With respect to genomic similarity, Hydrothermarchaeota MAGs discovered in BSmoChi-MAR share low values with MAGs extracted from Juan de Fuca Ridge ﬂank subsurface ﬂuids (SubFlu-JdFR) (Supplementary Figure S2), suggesting the wide phylogenetic distance within this phylum.

### Energy conservation metabolism

Components of type 3 and 4 [NiFe] hydrogenases (Hyc and Hyf) are patchily distributed in Hydrothermarchaeota MAGs; both of these hydrogenases are suggested to function together with formate dehydrogenase for the energy-conserving catalysis of formate to CO_2_ and H_2_ when the partial pressure of H_2_ is low^2^. Energy-converting hydrogenase A (EhaR only) and Energy-converting hydrogenase B (EhbQ only) are also found in all Hydrothermarchaeota MAGs; they are archaeal-specific membrane-associated energy-converting hydrogenases mainly found in hydrogenotrophic methanogens^3^. They catalyze the reversible reduction of ferredoxin by H_2_ oxidation which is driven by reverse electron transport^2^. Subsequently, the reduced ferredoxins could serve as electron donor and energy source in the fixation of CO_2_ to Formyl-MFR in THMPT-WL pathway^2^. The subsequently oxidized ferredoxins could also serve as electron acceptors in the converting of 2-Oxo acid to acetyl/succinyl-CoA and acetate by a series of ferredoxin oxidoreductase (e.g., Por, Kor, Ior, and Aor); then, the pool of reduced ferredoxins is replenished afterward. The Complex I-IV of electron transfer phosphorylation and V/A-type H^+^/Na^+^ transporting ATPase are found in all MAGs, responsible for chemiosmotic energy-conservation, transferring electrons of NADHs generated from central carbon metabolism to terminal electron acceptors and producing ATPs^4^.

**Supplementary Figures**

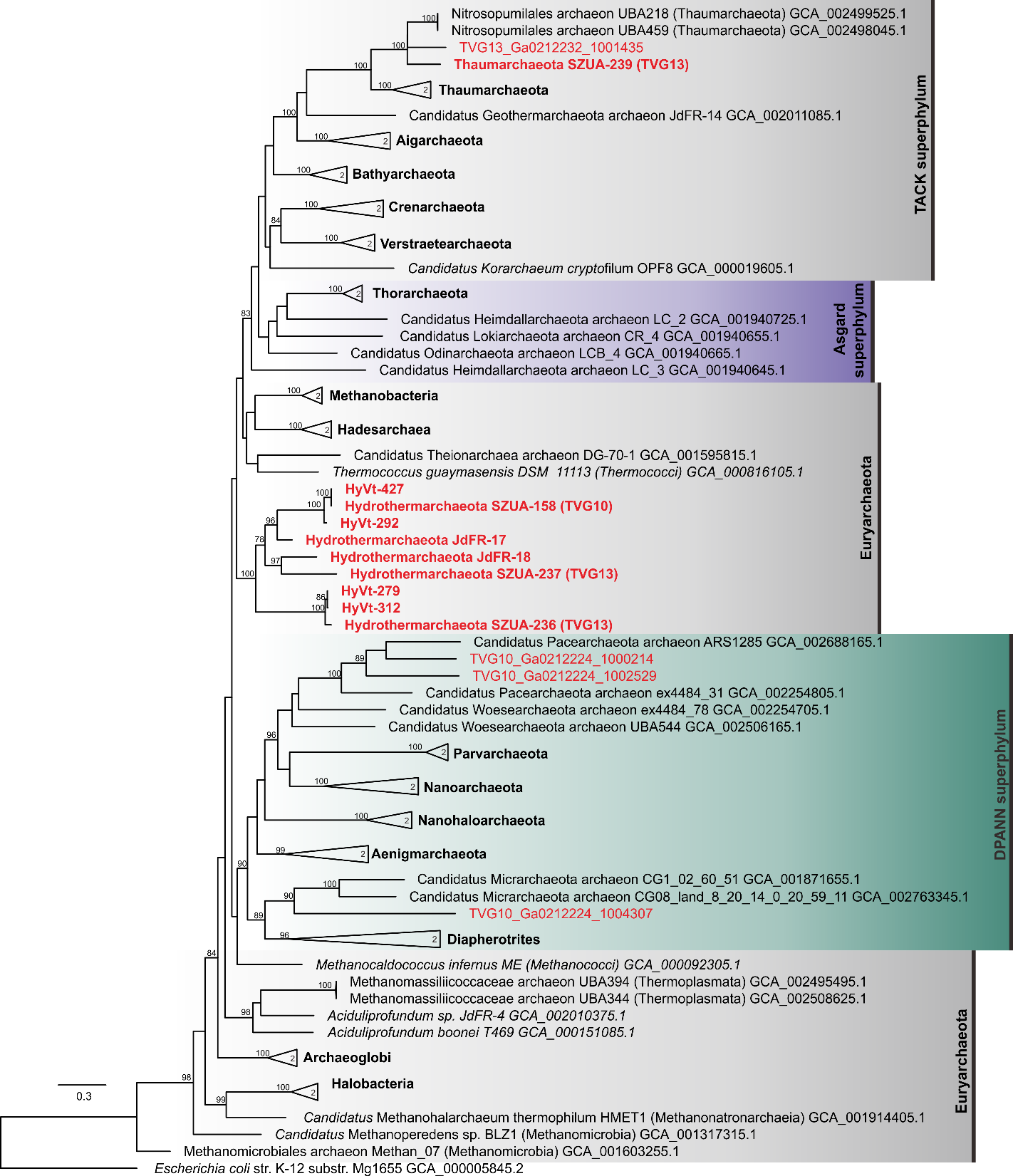
**Figure S1.** Phylogenetic tree of assembled archaeal MAGs and scaffolds based on 12 concatenated ribosomal proteins (L2, L3, L4, L5, L14, L16, L18, L22 and S3, S8, S17, S19 subunits). MAGs and scaffolds with less than 3 ribosomal proteins were pre-excluded. RAxML HPC v.8 was applied to reconstruct phylogeny with the best model as PROTGAMMAILG (suggested by ProtTest 3) and 100 times bootstrap iteration with autoMRE criterion. *Escherichia coli* K12 was used as the outgroup. Bootstrap supporting values over than 75% were labeled.

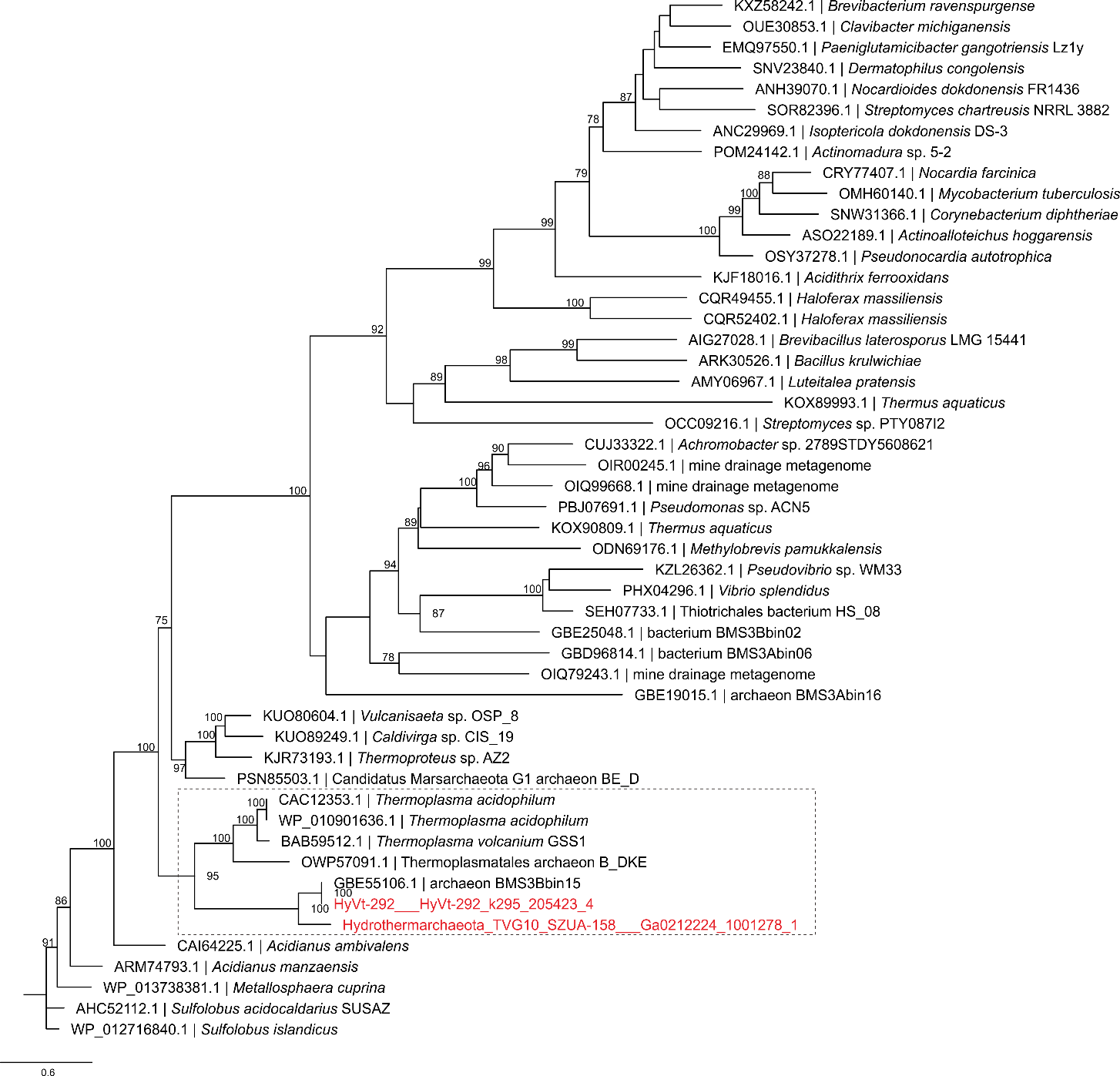
**Figure S2.** Phylogeny of AioB proteins. The protein alignment contains proteins from the NCBI database (May 16, 2018 updated, then, dereplicated at 60% similarity cutoff) and the best hits of AioB of this study by BLASTP in NCBI. This alignment is aligned by MAFFT^5^. This tree is constructed by IQ -TREE^6^ with settings of “-m MFP -mset LG,WAG -mrate E,I,G,I+G -mfreq FU -bb 1000”. The resulted ultrafast bootstrap values (UFBoot) are relatively higher than the corresponding default RAxML bootstrap values^7^. In this tree, only bootstrap values higher than 75% are labeled to the node. The sequences from Hydrothermarcheaota MAGs are labeled red. The UFBoot of the clade in dash line square is 95%, which means that of 95% possibility the topology of this clade is true.

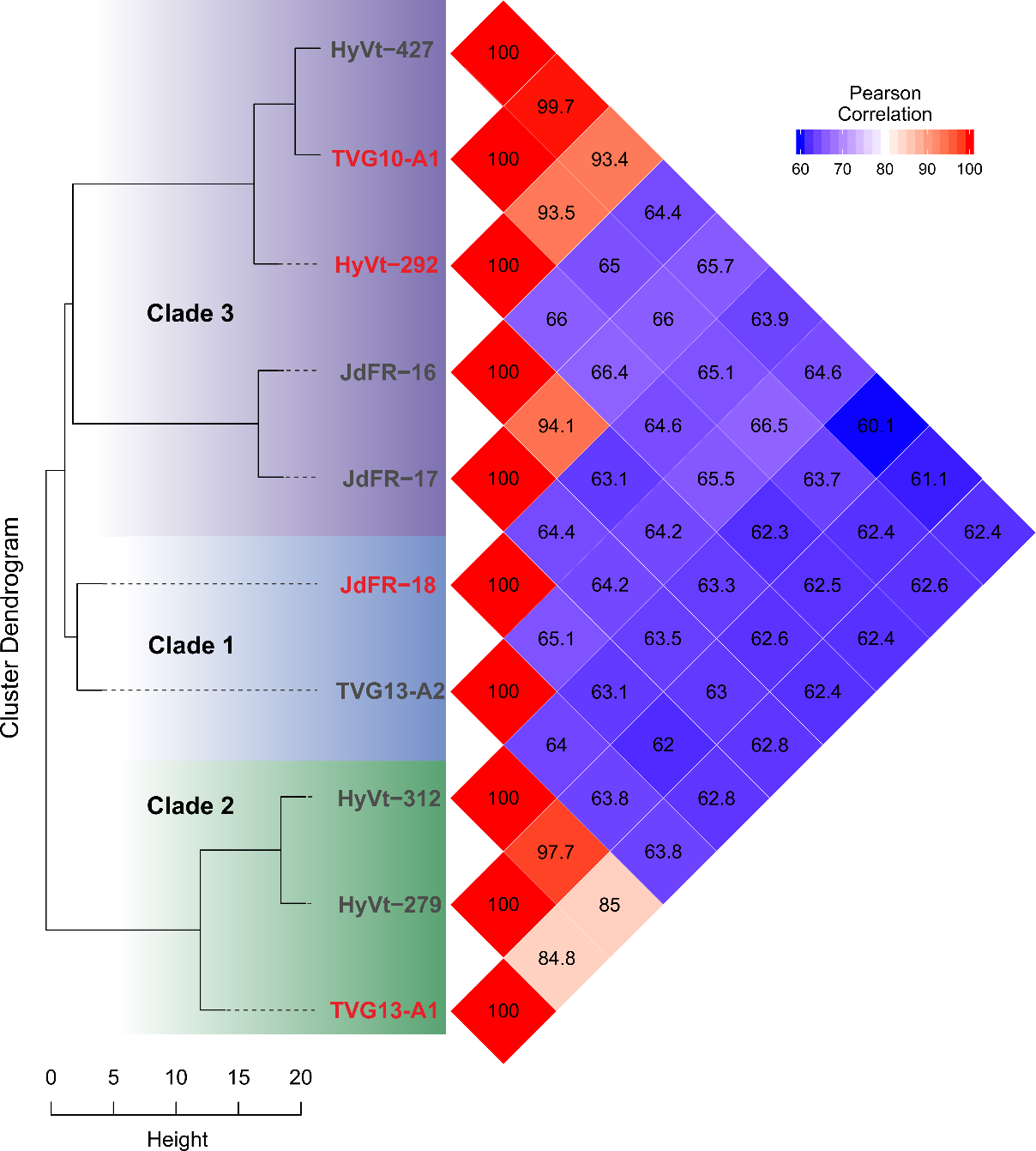
**Figure S3.** Pairwise orthoANI values among Hydrothermarchaeota MAGs. The matrix was reordered according to hierarchical clustering of the orthoANI values. Cluster dendrogram was generated from the matrix of orthoANI values. Red labeled MAGs are of high completeness (> 80%).

**Figure S4.** Comparative genomics of four Hydrothermarchaeota MAGs. The inner tree topology was assigned according to the matrix of COG functional categories and orthologous groups annotated by eggNOG-mapper. The items showed the presence/absence of gene clusters. The layers showed the presence/absence of specific functions (It is assigned according to the annotation results depicted in Figure 2. If there is one gene found in one specific function, we assign presence to this function).

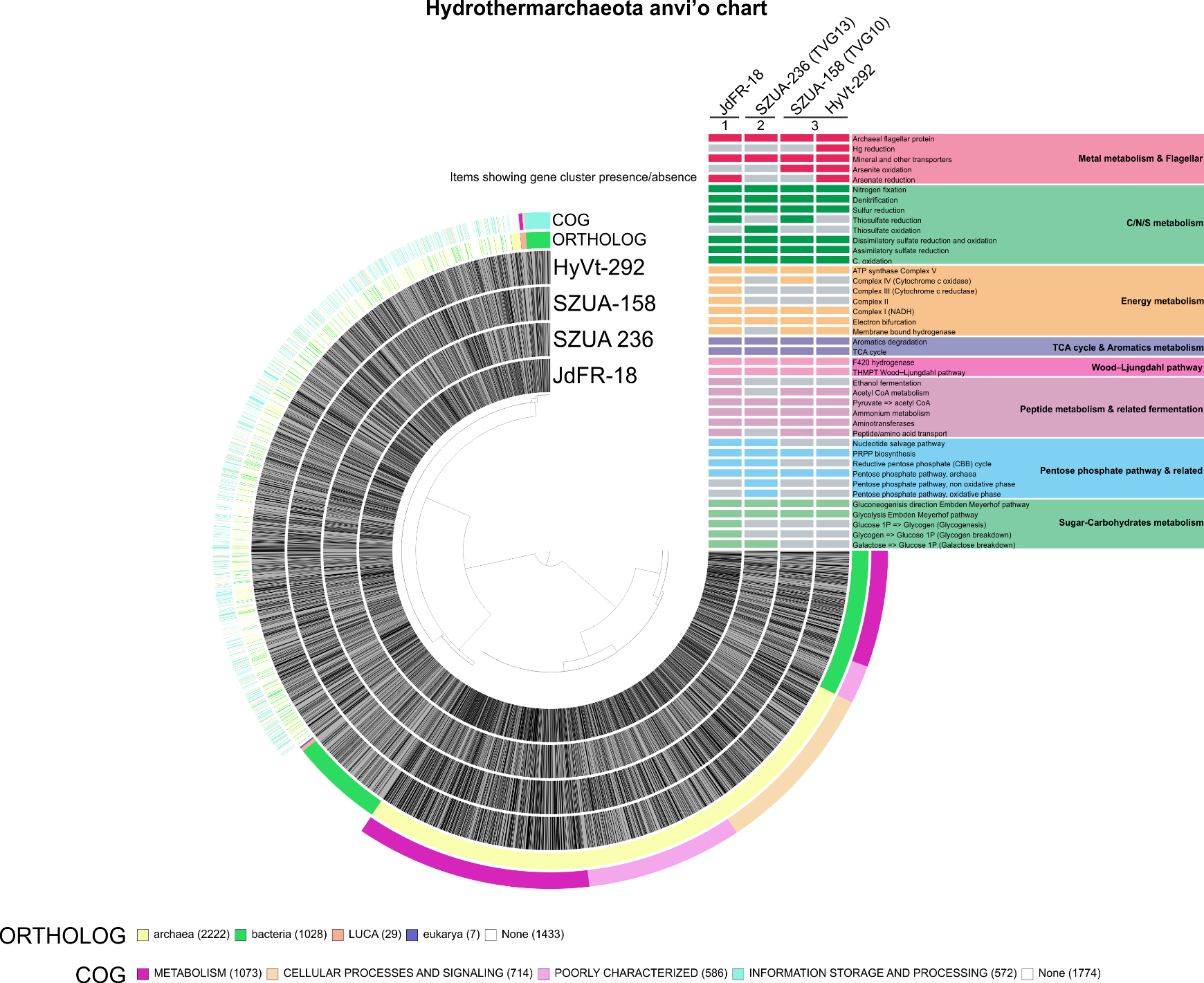

**Figure S5.** The phylogenomic tree of acquired 50 genomes for OG turnover event estimation. This phylogenomic tree is based on 12 concatenated ribosomal proteins (L2, L3, L4, L5, L14, L16, L18, L22 and S3, S8, S17, S19 subunits). This tree is constructed by IQ -TREE^6^ with settings of “-m MFP -mset LG,WAG -mrate E,I,G,I+G -mfreq FU -bb 1000”. The resulted ultrafast bootstrap values (UFBoot) are relatively higher than the corresponding default RAxML bootstrap values^7^. The tree is rooted by an outgroup of *Acidilobus saccharovorans* 345-15 (Crenarchaeota).

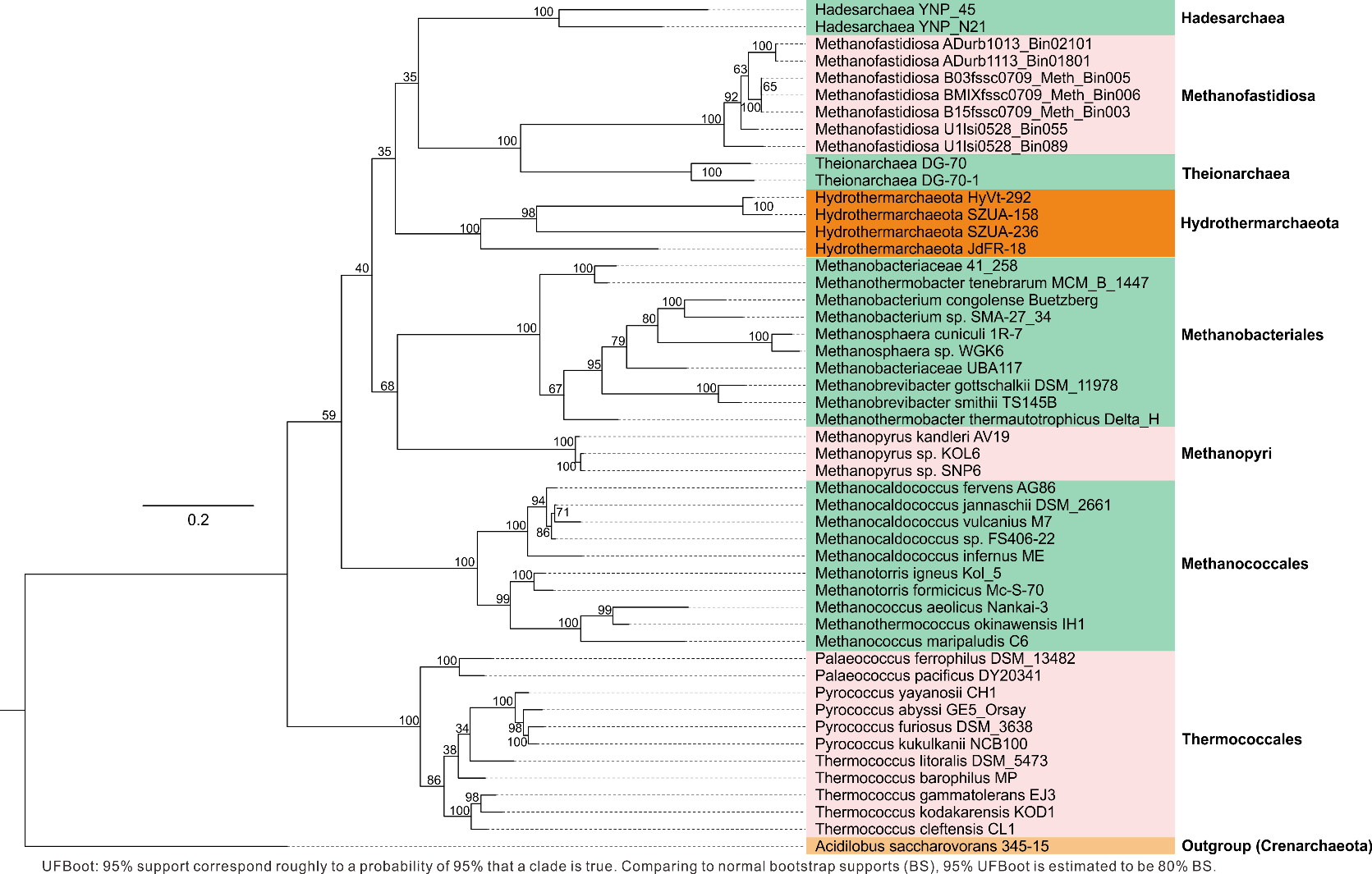

**Figure S6.** **a** Odd ratios of potential HGT genes to whole genomes for genes assigned as COG categories. The *p*-value and odd ratio were calculated by Fisher's Exact Test. The *p*-values indicate significant deviations between HGT genes and whole genomes (with < 0.05 as “*”, < 0.01 as “**” and < 0.001 as “***”). **b** The distribution of COG categories of potential HGT genes.
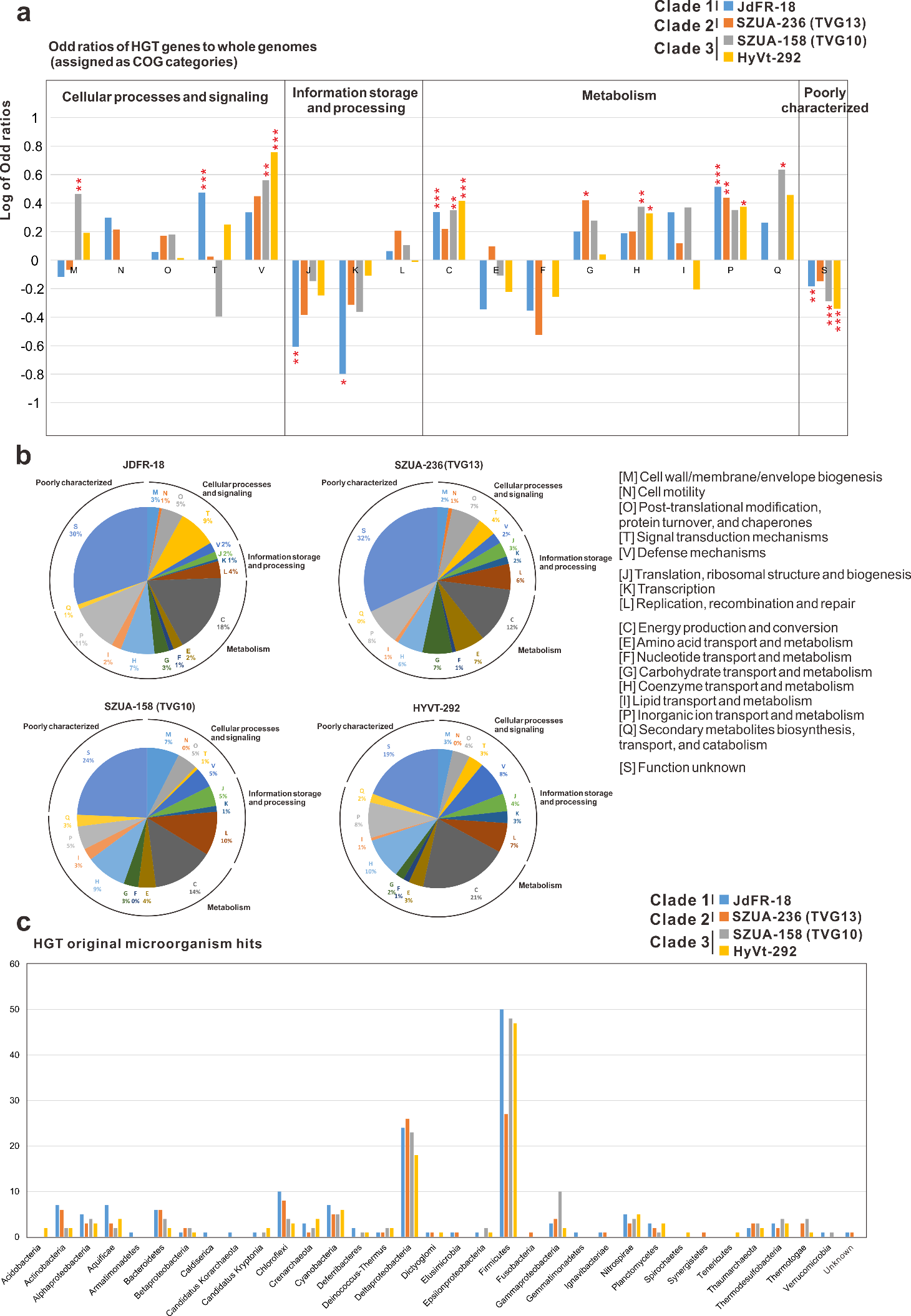
**c** The distribution of the original microorganisms of the potential HGT hits. Microorganisms were divided into phylum level.

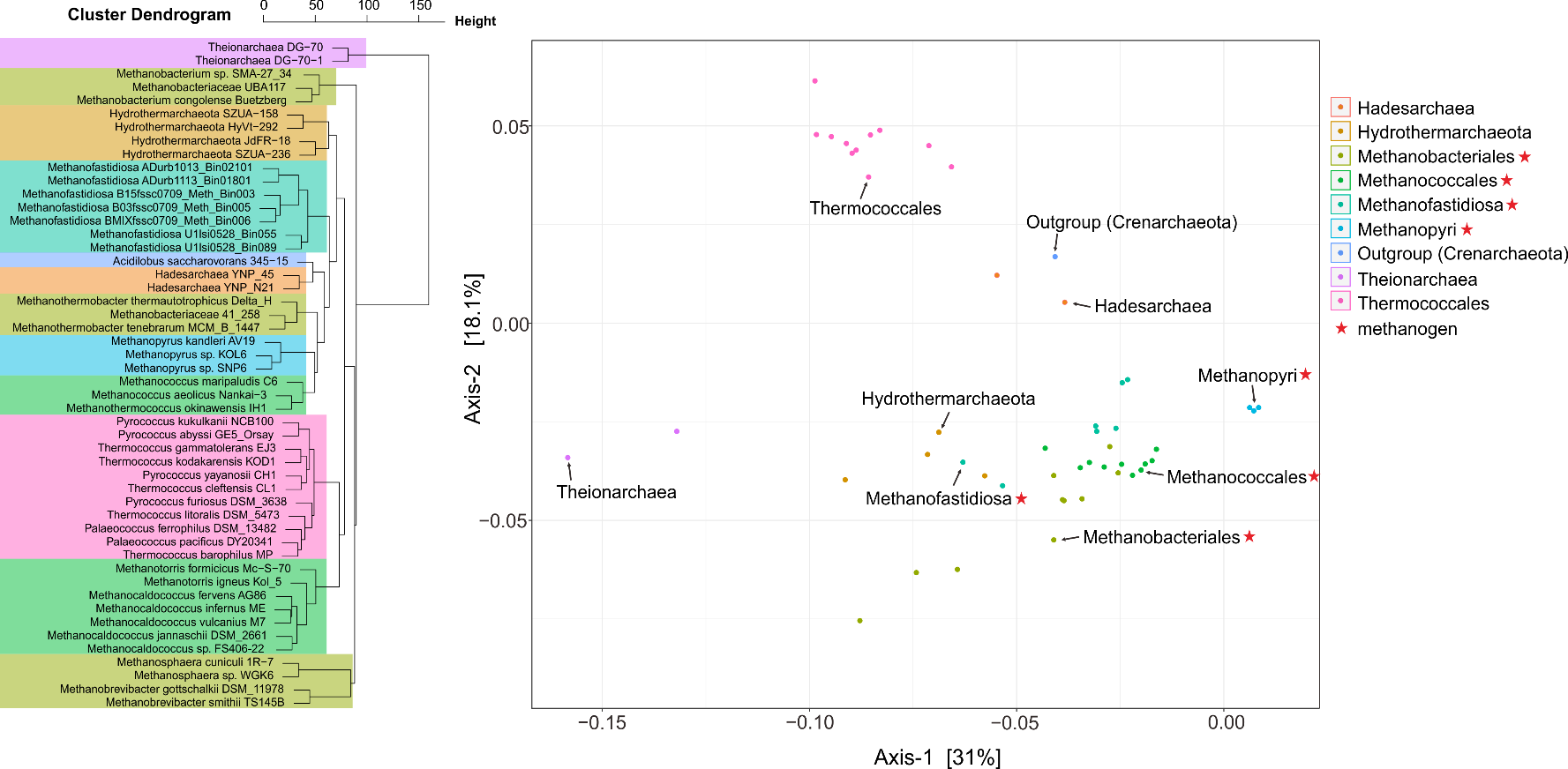
**Figure S7.** The PCoA and cluster dendrogram figures of 50 genomes for evolutionary analysis based on the dissimilarity matrix calculated from the OG component table. The PCoA ordination is based on Bray-Curtis distance calculating method. All of the 6750 inferred OGs shared among 50 genomes are included.
