## Supplementary material for "Genome and community-level interaction insights on wide carbon utilizing and element cycling function of Hydrothermarchaeota from hydrothermal sediment": Table S4.Proposed taxonomic level.docx

**Supplementary Table S4.** Proposed taxonomic level based on sequence identity range

| **Taxonomic group** | **Alternative name** | **Median sequence identity^a^** | **Median sequence identity^b^** | **Proposed taxonomic level based on sequence identity range^c^** |
| --- | --- | --- | --- | --- |
| Hydrothermarchaeota | Marine Benthic Group E (MBG-E) | 83.9 | 80.8 | Phylum |

The 16S rRNA gene diversity for assigning taxonomic level was conducted by picking representative sequences by QIIME [1] and being subjected to pairwise sequence identity analysis by BioEdit [2]. The median values of pairwise sequence identities were used.

a. Median sequence identity of representative sequences with 0.97 similarity cutoff from the SILVA database with sequence length over 1400 bp and pintail value over 75 (Nov 22, 2017 updated SSU Ref 128).

b. Median sequence identity of representative sequences with 0.97 similarity cutoff from the SILVA database with sequence length over 1200 bp and pintail value over 75 (Nov 22, 2017 updated SSU Ref 128).

c. This is according to the statistical report from the reference [3].

1.	Caporaso JG, Kuczynski J, Stombaugh J, Bittinger K, Bushman FD, Costello EK, Fierer N, Pena AG, Goodrich JK, Gordon JI *et al*: QIIME allows analysis of high-throughput community sequencing data. Nat Methods. 2010;7:335-336.

2.	Hall T: BioEdit: an important software for molecular biology. GERF Bull Biosci. 2011;2:60-61.

3.	Yarza P, Yilmaz P, Pruesse E, Gloeckner FO, Ludwig W, Schleifer K-H, Whitman WB, Euzeby J, Amann R, Rossello-Mora R: Uniting the classification of cultured and uncultured bacteria and archaea using 16S rRNA gene sequences. Nat Rev Microbiol. 2014;12:635-645.
